# DIMA: Data-driven selection of a suitable imputation algorithm

**DOI:** 10.1101/2020.10.13.323618

**Authors:** Janine Egert, Bettina Warscheid, Clemens Kreutz

## Abstract

**Motivation:** Imputation is a prominent strategy when dealing with missing values (MVs) in proteomics data analysis pipelines. However, the performance of different imputation methods is difficult to assess and varies strongly depending on data characteristics. To overcome this issue, we present the concept of a *data-driven selection of a suitable imputation algorithm* (DIMA).

**Results:** The performance and broad applicability of DIMA is demonstrated on 121 quantitative proteomics data sets from the PRIDE database and on simulated data consisting of 5 – 50% MVs with different proportions of missing not at random and missing completely at random values. DIMA reliably suggests a high-performing imputation algorithm which is always among the three best algorithms and results in a root mean square error difference (ΔRMSE) ≤ 10% in 84% of the cases.

**Availability and Implementation:** Source code is freely available for download at github.com/clemenskreutz/OmicsData.

## 1 Introduction

In many scientific fields, incomplete data still raises questions with regard to their analysis and interpretation. When dealing with missing values (MVs) many researchers are not aware of the serious impact imputation - a crucial step in many analysis pipelines - or its omission may have on the statistical power of downstream analyses or on the estimation bias [Janssen et al., 2010, Välikangas et al., 2017, Wang et al., 2017]. There are many computational approaches that cannot deal with missing data. For such methods, imputation is mandatory in order to avoid restrictions to a small number of proteins that could be observed in all samples. However, the choice of a high-performing algorithm is rather ambiguous because the origin, e.g. technical or biological constraints, and the distribution of missing data are often unknown. The statistical categories: Missing At Random (MAR), Missing Not At Random (MNAR) or Missing Completely At Random (MCAR) [Rubin, 1976] refer to the origin of missing data and are widely used in the analysis of MVs [Karpievitch et al., 2012, Lazar et al., 2016, Wei et al., 2018]. Although there exists Little’s test to determine if the MVs are MCAR or MNAR [Little, 1988], the relation and distribution of MNAR and MCAR values can not be determined.

Imputation still gives rise to lively discussions in the bioinformatics field and within the last few years many papers have been published concerning the application and performance of various imputation algorithms. In one of these publications, eight imputation methods are examined on microarray expression profiles, of which *least squares adaptive* (*LSA*), *local least squares* (*LLS*) and *Bayesian principal component analysis* (*BPCA*) perform best [Brock et al., 2008]. To determine the chemical rank based on the *Toxicological Priority Index*, *Mean* imputation has been proven to be the algorithm of choice [To et al., 2018]. In contrast, when imputing spatially explicit plant traits, *multivariate imputation using chained equations* (*MICE*) performs best [Poyatos et al., 2018]. Hence, the performance of different imputation algorithms highly varies depending on data acquisition and data characteristics [de Souto et al., 2015, Liu and Dongre, 2020, Rodwell et al., 2014]. Therefore, making a general suggestion for an optimal imputation strategy seems to be unfeasible [Brock et al., 2008, Audoux et al., 2017, Webb-Robertson et al., 2015]. The impact of imputation on data analysis pipelines by investigating missing data occurrence patterns has been introduced in [Hrydziuszko and Viant, 2012]. DIMA extends this approach and learns the probability of MV occurrences depending on the protein, sample and mean protein intensity by logistic regression model. The regression coefficients are then employed to imitate a likewise MV distribution and therewith the performance of multiple imputation algorithms can be examined. With DIMA, a valid recommendation of a high-performing imputation algorithm specifically tailored to the user-supplied data set and its MV distribution is offered.

## 2 Methods

### 2.1 Illustration data

To illustrate the concept of the data-driven selection of a suitable imputation algorithm, DIMA is applied to a liquid chromatography tandem mass spectrometry (LC-MS/MS) data set with complete native stable isotope labeling by amino acids for analyzing quantitative changes in the proteome of mitochondrial gene deletion strains in comparison to wild-type cells [Dannenmaier, 2018]. This data set (PXD010704) is generated from mitochondria-enriched fractions of Saccharomyces cerevisiae wild-type, *sdh5*Δ, *phb1*Δ and *coi1*Δ cells, in which 1,609 proteins were identified with 56% MVs. The data set is depicted in Figure 1 as original data *O*.

**Figure 1:**
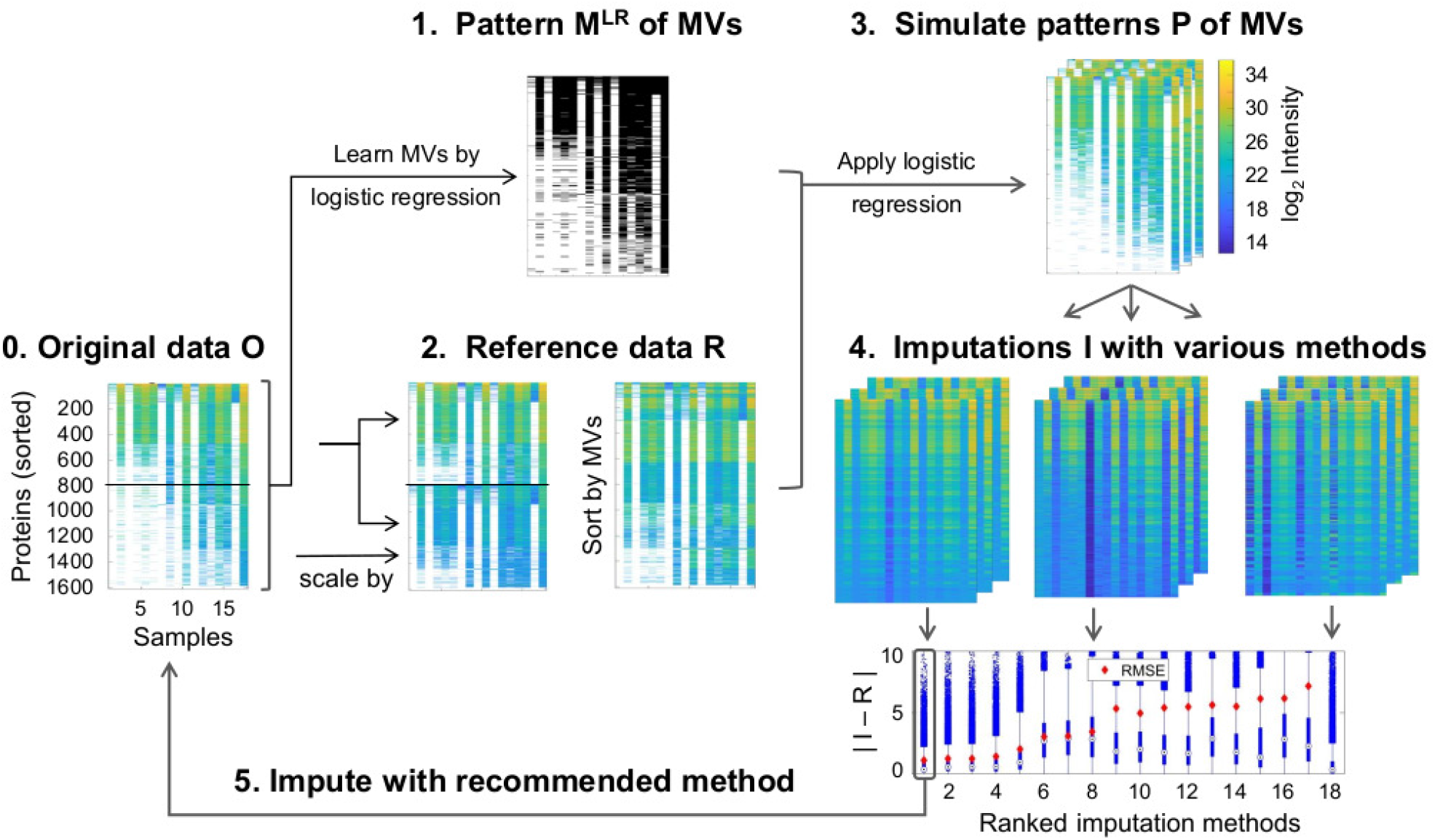
DIMA analysis pipeline, exemplarily applied to an illustration data set [Dannenmaier, 2018]. **0.** The original data *O* with rows sorted according to the number of MVs and mean protein intensity. **1.** The pattern *M^LR^* of MVs is learned by logistic regression using the protein and sample as factorial predictors, plus the mean protein intensity. **2.** A data subset *R* with few MVs is defined. **3.** Patterns *P* of MVs are simulated by the logistic regression model and the respective coefficients of step 1 and incorporated in the reference data *R*. **4.** Multiple imputation algorithms are applied to the reference data *R* with incorporated MVs and the RMSE is calculated. For visualization purposes the boxplot is limited to [0, 10]. **5.** The best-performing imputation algorithm on the reference data R is recommended for the original data *O* and imputation of *O* is conducted.

### 2.2 PRIDE data

To show its broad applicability and performance, DIMA is applied to 121 publicly available data sets from the PRIDE database, which are acquired by downloading all files from the ftp-server “ftp.PRIDE.ebi.ac.uk” containing the search pattern “MaxQuant” or “proteinGroups” and file extensions .txt, .xlsx, .csv, .tar, .tar.gz, .zip or .rar from 05 – 07, 2020. The 121 PRIDE data sets comprise [190 – 13,430] proteins, [2 – 240] samples and [2.6 – 94.3]%MVs, with a mean number of (3100 ± 2500) proteins, (19 ± 23) samples and (41 ± 26)% MVs. 35% of the data sets contain more than 50%MVs.

### 2.3 Simulation study

In addition to experimental data sets, a simulation study is performed to analyze the influence of randomness, i.e. the ratio of MCAR and MNAR values and the frequency of MVs, on the imputation performance. The protein intensities

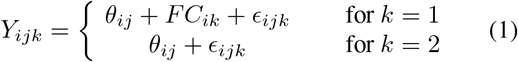

are calculated with the baseline average *θ_ij_* ~ *N*(18.5, *IG*(1,1)) of peptide *j* matched to protein *i* and with a noise term *ϵ_ijk_* ~ *N*(0, *IG*(2,1)). For condition *k* =1 a fold change *FC_ik_* ~ *N*(0, *IG*(1.5,1)) is added to the protein intensity in order to represent differential expression. The variances are drawn from the inverse gamma distribution *IG* [O’Brien et al., 2018].

To evaluate the impact of the frequency and randomness of MVs, various combinations of MVs and MNAR/MCAR proportions are incorporated in the simulated data *Y*. The MCAR values are set randomly. As suggested in [O’Brien et al., 2018], MNAR values are drawn from

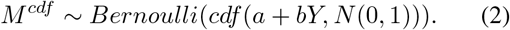

with *M^cdf^* ∈ {0, 1} with an intensity dependent probability. *M^cdf^* = 1 indicates available observations and *M^cdf^* = 0 indicates missing values. The cumulative distribution function is calculated with the rate *b* of MNAR values and the b-quantile *a* = *Q_b_*(*Y*) of the protein intensities *Y*. The simulated data set *S* with incorporated MVs consists of the simulated intensities *Y* for *X* = 1 and MV incorporations for *X* = 0. For each MV and MNAR combination, 300 data sets with 500 proteins and 20 samples which consist of two conditions *k* = {0, 1} and ten replicates are simulated.

### 2.4 DIMA

DIMA assesses and suggests imputation algorithms for a user-defined data set *O*. The method consists of five main steps which are depicted in Figure 1 and which are described in more detail in the following subsections:

1. The pattern *M^LR^* of MVs in the original data *O* is learned by logistic regression analysis (Section 2.5).
2. A reference data *R* with fewer MVs is constructed from the original data *O* to evaluate imputation performance on.
3. To generate a pattern of missing data with a similar distribution as in the original data, the logistic regression model of step 1 is applied to the reference data *R* (Section 2.6). Bernoulli trials are performed to simulate different patterns *P* of MVs.
4. Various imputation algorithms (Section 2.7) are applied to the reference data *R* with patterns *P* of MVs and ranked by their root mean square error (RMSE). The best-performing algorithm is given by the lowest average 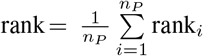 over all pattern simulations (Section 2.8).
5. The best-performing imputation algorithm of step 4 is recommended as imputation algorithm for the original data set *O* and imputation of *O* is performed.

### 2.5 Learn pattern of missing values

The pattern *M^LR^* of missing values is defined as:

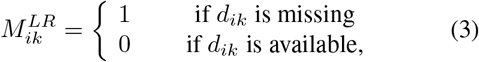

with *d_ik_* corresponding to the data value for protein *i* within sample *k*. The probability

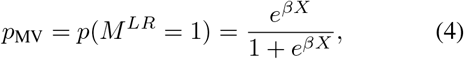

of a specific data point being missing is described with a logistic regression model. The columns of the design matrix *X* represent data-characteristic predictor variables, including the mean protein intensity and the protein and sample as factorial predictors. In addition, DIMA offers optional predictors such as protein ratios, molecular weight or quality scores if included in the input data file.

The predictors are standardized [Marquardt, 1980, Menard, 2011] and a weak regularization of the regression coefficients *β* for the factorial predictors has been performed to decrease the variance of the estimated parameters and to prevent non-identifiability [Kreutz, 2016]. For large data sets (> 1000 proteins) random subsampling of the proteins is performed to decrease computational effort. Each protein is drawn once and the regression coefficients *β* are set to the sampling means.

### 2.6 Simulate patterns of missing values

The reference data set *R* should contain few MVs and should have similar intensity distribution as the original data set *O*. For DIMA, *R* is constructed by selecting a proportion *c* of proteins with the least number of MVs. This subset is then augmented by the same *c* proteins and scaled to the mean and standard deviation of the remaining proteins. By default, the fraction of proteins with the least number of MVs is set to *c* = 0.5. To reliably recommend an imputation strategy, a pattern *P* of MVs similar to the original pattern *M^LR^* is simulated. The probability *p*_MV_ of MV occurrence is calculated with the coefficients *β* and equation 4 and Bernoulli trials are performed. Depending on the size of the input data *n_P_* =5 – 20 patterns are generated.

### 2.7 Imputation algorithms

By applying DIMA, 30 imputation algorithms from 13 R-packages which are based on a wide variety of imputation strategies such as expectation-maximization (EM), singular value decomposition (SVD), principal component analysis (PCA), neural network or sequential imputation are evaluated. The algorithms are depicted in Table S1 with their algorithm name, package name, required input parameters, explanations and references.

### 2.8 Ranking of imputation algorithms

To compare the performance of the different imputation algorithms, the root mean square error

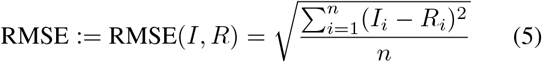

between the measured protein intensities *R_i_* of the reference data set and the imputed protein intensities *I_i_* is calculated. Here, *n* denotes the total number of imputed entries.

For each generated pattern the imputation algorithms are ranked by their RMSE. The approach with the lowest mean rank over all pattern simulations is recommended as best-performing algorithm for the original data *O*. If for any reason an imputation algorithm fails, the highest rank is assigned for this respective imputation. Therewith, the algorithm may still be recommended by DIMA, even though this is unlikely. An imputed data set which still contains MVs is by definition treated as an imputation failure.

A minimal RMSE indicates that the imputation is on average closest to the true values, which is intended in most cases for subsequent analysis steps, like performing fold changes or cluster analysis. For statistical tests, however, also the variability of a data set is substantial and therefore the imputation has to reflect the same distribution and variance as the underlying truth. Therefore, alternative criteria can be employed for the ranking of imputation algorithms. For the t-test, the RMSE*_t_* := RMSE(*t_R_*, *t_I_*) serves as rank criterion where *t* is the t-test statistics calculated from the observed data *R* and accordingly from the imputed data *I*. The associated sample indices for the underlying null hypothesis have to be specified by the user.

To verify variations in the imputed intensity distributions, the differences in variances for the observed data *R* and imputed data *I* are examined with a two-sample F-test. Here, the p-value

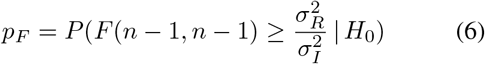

under the null hypothesis *H*_0_ of equal variances 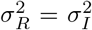 serves as criterion.

## 3 Implementation

DIMA is implemented in Matlab and publicly available at github.com/clemenskreutz/OmicsData. The analyses were performed on an Intel Xeon E5-2640v3 CPU with 16 cores on the BwForCluster MLS&WISO Development.

A proteomics data object is created by

~~~
O = OmicsData(file);
~~~

where .xls, .txt and .mat files as well as a numeric input are accepted, e.g. the MaxQuant output tables can serve as file inputs here. DIMA is executed via

~~~
O = DIMA(O,[algorithms],[bio]);
~~~

The imputation algorithms are taken according to Table S1 or alternatively can be specified by the user. A fast version, which only runs the nine most frequently recommended algorithms based on the 121 PRIDE data sets, is also implemented and available by setting algorithms = *’fast’*. The optional third input argument is a flag if biological information such as protein ratios, molecular weight or scores should be taken into account. After applying DIMA the suggested algorithm and the respective imputation are stored in the proteomics data object and can be accessed via:

~~~
algorithm = get(O,’DIMA’);
data = get(O,’data’);
~~~

## 4 Results

In the following, DIMA is first demonstrated on an illustration LC-MS/MS data set (see section 2.1). To show its broad range of applicability, DIMA is applied to 121 PRIDE data sets and furthermore to simulated MNAR and MCAR data sets to evaluate its performance.

### 4.1 Illustration data

The application of DIMA to the LC-MS/MS illustration data set is depicted in Figure 1. Imputation performance on the simulated patterns *P* of MVs is shown in step 4 as boxplots. The three best algorithms show an equally good performance with a low RMSE and a low absolute error between the imputed data *I* and the reference data *R*. The performance measures RMSE, RMSE*_t_* and *p_F_* with their respective ranking are depicted in Table 1 for the five bestperforming and the least-performing imputation algorithms. By default, the lowest mean rank of the RMSE over all pattern simulations *P* is taken for recommendation of an imputation algorithm.

**Table 1:** The RMSE of the imputed data values, RMSE of the t-test statistics, p-value of the F-test statistics and the respective rankings are illustrated for the three best-performing and the least-performing imputation algorithms on the illustration data set. In bold the best-performing algorithm for the respective criterion.

| algorithm | RMSE | $\overline{\text{rank}}$ | RMSE<sub>t</sub> | $\overline{\text{rank}}_t$ | $p_F$ | $\overline{\text{rank}}_F$ | time [min] |
| --- | --- | --- | --- | --- | --- | --- | --- |
| missForest | **0.017** | **1.0** | 2.7 | 3.3 | 0.186 | 2.5 | 11.5 |
| MIPCA | 0.020 | 2.3 | 3.0 | 6.8 | 0.002 | 4.1 | 1.6 |
| imputePCA | 0.020 | 2.8 | 2.9 | 5.6 | **0.364** | **1.7** | 2.3 |
| kNNImpute | 0.023 | 4.0 | **2.5** | **1.4** | 0.184 | 2.4 | 0.8 |
| SVDImpute | 0.033 | 5.1 | 2.7 | 2.8 | <1e-10 | 6.4 | 0.1 |
| ... | ... | ... | ... | ... | ... | ... | ... |
| QRILC | 0.516 | 20.3 | 3.8 | 14.8 | <1e-10 | 17.4 | 0.3 |

The best-performing algorithm for this exemplary data set is the random forest algorithm from the R package *missForest* [Stekhoven and Buhlmann, 2012] with an average rank = 1 over all pattern simulations, followed by the PCA algorithms *MIPCA* and *imputePCA* from the R package *missMDA*. The other imputation algorithms result in higher values for RMSE. The computation time for each imputation algorithm ranges from 6 s to 11.5 min. Applying DIMA to the illustration data set takes 28 min in total. The null hypothesis of equal variances in the original compared to the imputed data set is rejected (p_F_ < 0.01) for 17 and not rejected for 4 of the 18 imputation algorithms. To determine if the protein intensities obtained from the deletion strains differ significantly compared to the wild-type cells, t-test statistics is applied. The RMSE of the t-test statistics over all identified proteins is smallest for the imputation algorithm *kNNImpute*. Here, the sequential algorithm which performs worst in representing the true data values performs best in representing the t-test statistics of equal means in the different cell types.

Different rankings are obtained for the imputation algorithms depending on the rank criterion (Table 1). Therefore, the selection of the rank criterion is a crucial step and has to be made with regard to the downstream analysis steps.

If the original data *O* includes proteins without a data value for any sample i.e. a row with complete missingness, the user could consider removing those proteins from the data set *O* beforehand. Within DIMA, proteins with complete missingness are included in the patterns *P* of MVs and the evaluation of the imputation algorithms. However, the R-packages *pcaMethods* and *impute* fail if there are proteins without intensities.

### 4.2 PRIDE data

DIMA is applied to 121 data sets from the PRIDE database (see section 2) to demonstrate its general applicability and to evaluate the various imputation algorithms on multiple data sets. For 44% of the data sets DIMA suggests the robust sequential algorithm *impSeqRob* and for 26% of the data sets DIMA recommends the sequential algorithm *impSeq* (Figure 2). Both approaches are available in the *rrcovNA* package and are based on sequentially imputing each MV by minimizing the determinant of the covariance matrix. Further suggested imputation algorithms are from the R packages *missForest* (12%), *missMDA* (13%), *pcaMethods* (4%) and *imputation* (2%). On average, DIMA takes (2.1 ± 1.8) minutes cputime per data set, with a median of 1.6 minutes.

**Figure 2:**
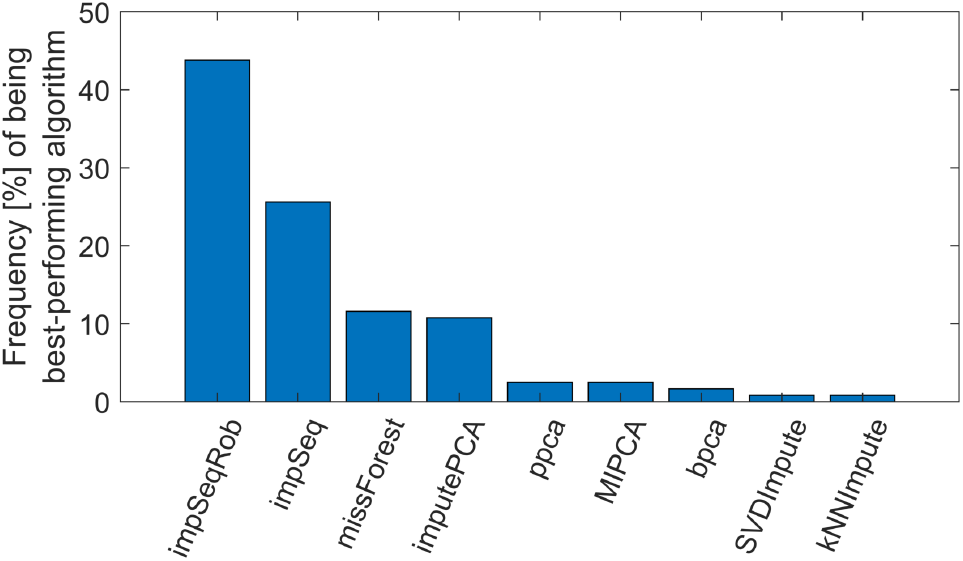
DIMA is applied and evaluated on 121 Pride data sets. Nine algorithms compete for being recommended as best-performing algorithm. The R-package *rrcovNA* with its algorithms *impSeqRob* and *impSeq* is selected in 69% of the Pride data sets, *missForest* in 12%, *imputePCA* in 11% and for 8% of the Pride data sets another algorithm is suggested.

### 4.3 Simulation study

To evaluate whether DIMA is capable to reliably identify high-performing imputation approaches, a data simulation study is conducted. The data is simulated as described in section 2.3 with MV ∈ [5, 50]% and MNAR ∈ [0, 100]%. Furthermore, the simulated data sets *S* with incorporated MVs are imputed and assessed by comparing it to the simulated data *Y* before MV incorporation. This assessment and the respective ranking of the methods is termed *direct imputation assessment* in the following, and serves as the assumed true algorithm ranking to evaluate the performance of DIMA.

In Figure 3, the average rank of the recommended algorithm by DIMA achieved in direct imputation assessment for the respective MV/MNAR combination and over 300 data simulations is shown as color map. The average rank ranges from 1.0 to 2.7 indicating that DIMA recommends one of the three best algorithms in all cases independent of the number of MV or MNAR values. In each box, the mean RMSE difference 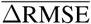 (first entry) of the algorithm recommended by DIMA (second entry) compared to the best-performing algorithm with direct imputation assessment (third entry) is averaged over all simulated data sets S and the respective MV/MNAR combinations. Here, the most frequent algorithm recommendation is illustrated. For MV < 20% the two methods *regression* and *pmm* of the algorithm *aregImpute* from the R package *Hmisc* are most frequently recommended by DIMA and by direct imputation assessment. Both methods apply a flexible additive regression model to bootstrap samples.

**Figure 3:**
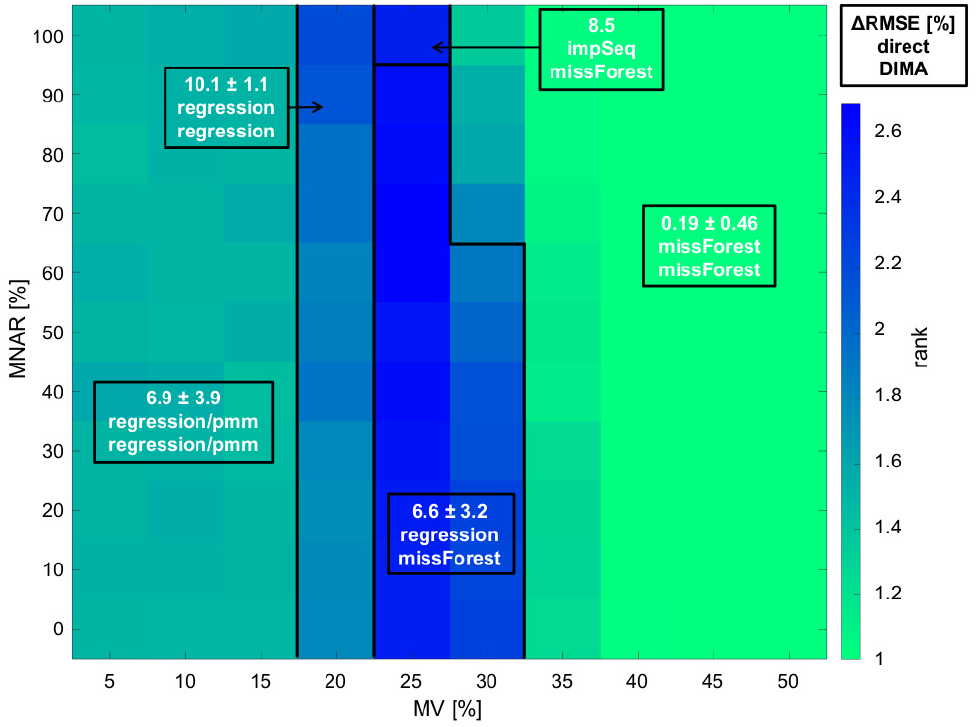
The performance of DIMA is evaluated on simulated data *S* with incorporation of various proportions of MV and MNAR/MCAR values. The best-performing imputation algorithm recommended by DIMA is then compared to the rank and RMSE obtained by direct imputation assessment. The average rank over 300 data simulations is shown as color map. The algorithm recommended by DIMA is within the top three approaches in all cases. The 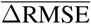 (first entry) of the recommended algorithm (second entry) compared to the best-performing algorithm by direct imputation assessment (third entry) over all simulated data sets and over the respective MV/MNAR combinations is 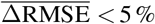 in 62 % of all cases.

For MV > 30% the random forest algorithm *missForest* shows best imputation performance. In between for 20% ≤ MV ≤ 30% the two R packages *missForest* and *Hmisc* compete as best-performing imputation algorithm depending on the simulated data set which results in greater rank and RMSE difference. In 84% of the cases 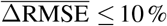.

A second data simulation adapted from [Lazar et al., 2016] is performed and the performance for the different MV/MNAR combinations is shown in Figure S2. For these simulated data sets, the approach suggested by DIMA is within the top three approaches in 78% of the cases and results in 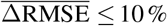 in 80% of the cases. Yet, data sets with MNAR > 90% or MV < 10% result in greater deviations. Comparing the imputation performance for the two data simulation processes, the recommended algorithms differ strongly. However, the low ranks of the recommended algorithms elucidate that DIMA indeed learns the respective data characteristics. This emphasizes the importance of learning the individual data structure to decide for a high-performing imputation algorithm.

## 5 Discussion

In recent years, many papers have been published concerning imputation strategies, because downstream analyses are highly affected by this processing step. To facilitate the decision for a well-performing imputation method, a novel concept for data-driven selection of a suitable imputation algorithm is presented. DIMA has the advantage that it combines the learning of the individual pattern of MVs in a given data set with the testing of many different imputation strategies to reveal the best-performing algorithm for the specific input data.

We were able to demonstrate that DIMA succeeds in recommending a high-performing imputation algorithm for different combinations of MV frequencies and MNAR/MCAR ratios. The suggested imputation algorithms are within the top three methods for all combinations. However, in just 60% of the cases the best-performing algorithm was correctly identified. This is due to a comparably strong imputation performance of several methods, so that the ranking within these methods highly depends on the data realization.

Similar MV and MNAR ratios lead to similar algorithm recommendations (Figure 3). However, the algorithm recommendations between the two considered simulation approaches differ strongly (Figure S2). Simulating different data characteristics by [O’Brien et al., 2018] and by [Lazar et al., 2016] show the importance of selecting a suitable imputation strategy for the respective data properties.

To show its broad applicability, DIMA was applied to more than 100 data sets of the PRIDE database. The algorithm *impSeqRob* from the R package *rrcovNA* most frequently performed best and could thus be used as a default imputation strategy.

A classification tree to predict a high-performing imputation algorithm based on data-specific properties such as number of proteins, number of samples, MVs, ratio of MNAR values, skewness or biological incidents like experiment type, instrument, species or post-translational modifications was implemented. However, we were not able to identify attributes which are predictive for selecting an optimal imputation algorithm.

A crucial aspect in choosing a suitable imputation algorithm is the subsequent downstream analysis, as the recommended algorithm may introduce bias. While some imputation methods strive to reflect the true value as close as possible, other algorithms focus on other data characteristics such as similar distributions or equal variances. Thus, DIMA provides several measures to evaluate target criteria such as closeness to the original value (RMSE), t-test statistics (RMSE_t_) or maintaining variances (*p_F_*).

DIMA currently comprises 30 imputation algorithms and takes (2.1 ± 1.8) minutes cputime per data set. Depending on the given data, variation of hyper-parameters for the individual imputation methods could result in a better performance. However, evaluation of hyper-parameters for all algorithms quickly leads to inflating computation time.

For large data sets and/or time saving purposes one could consider omitting the calculation of imputation algorithms which are not promising for good performance by prior knowledge e.g. algorithms which were not suggested by DIMA in the PRIDE data sets. A fast version of DIMA which only includes the nine most frequently recommended algorithms on the PRIDE data sets is implemented.

In this work, the high performance and effectiveness of DIMA is demonstrated on various quantitative mass spectrometry data sets. However, DIMA represents a general approach and thus can be applied to any data set regardless of its origin or scientific background. Particularly, data from other high-throughput techniques such as RNASeq or single-cell data would highly benefit from the recommendation of a high-performing imputation algorithm.

## 6 Acknowledgements

The authors acknowledge support by the state of Baden-Württemberg through bwHPC and the German Research Foundation (DFG) through grant INST 35/1134-1 FUGG. We gratefully thank Eva Brombacher, Lena Reimann, Wignand Mühlhäuser and Friedel Drepper for fruitful discussions about the topic.

