## supplement for "DIMA: Data-driven selection of a suitable imputation algorithm"

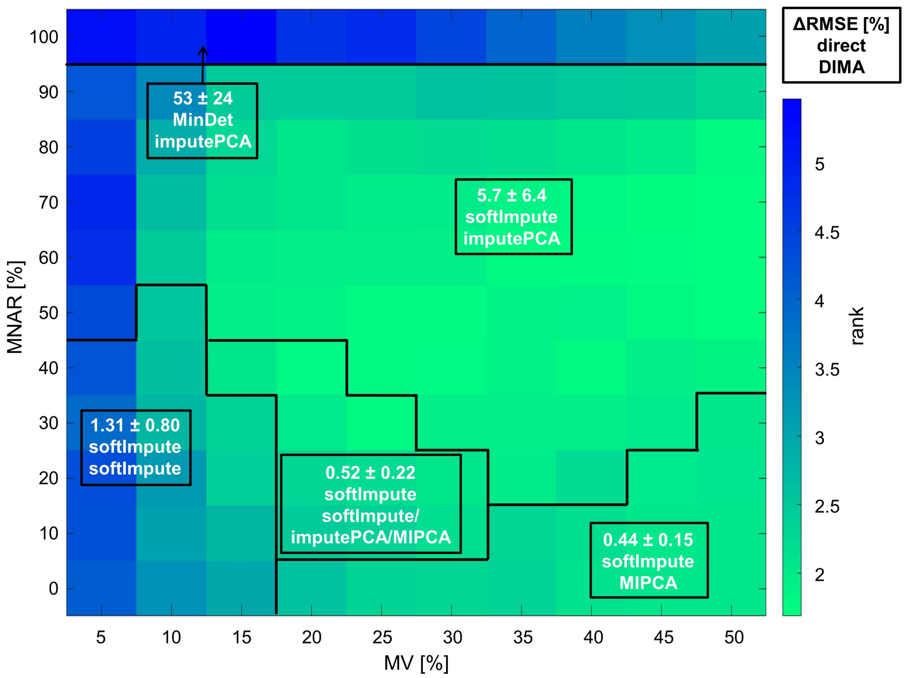


Figure 1: The performance of DIMA is evaluated on simulated data with incorporation of various proportions of MV and MNAR/MCAR values. The data simulation and MV incorporation is adapted from [Lazar et al., 2016]. In each box, the $\overline{\Delta \text{RMSE}}$ (first entry) of the recommended algorithm (second entry) compared to the best-performing algorithm by direct imputation assessment (third entry) over all simulated data sets is shown. The green color indicates a good performance of DIMA recommending a top three method in 91 % of the cases. In 80 % of the cases the $\overline{\Delta \text{RMSE}} \leq 10\%$.

| Algortihm | Explanation | R-package | Required input | Included in fast DIMA | References |
|---|---|---|---|---|---|
| amelia | bootstrap expectation maximization | Amelia | m = 1 | - | [Honaker et al., 2011] |
| aregImpute | additive regression with type pmm | Hmisc | n.impute = 1 | - | [van Buuren, 2012] |
| regression | aregImpute with type regression | Hmisc | n.impute = 1 | - | [van Buuren, 2012] |
| MinDet | deterministic minimal value | imputeLCMD | – | - | [Lazar et al., 2016] |
| MinProb | probabilistic minimal value | imputeLCMD | - | - | [Lazar et al., 2016] |
| QRILC | quantile regression | imputeLCMD | - | - | [Lazar et al., 2016] |
| kNNImpute | k-nearest neighbor | imputation | k = 3 | yes | [Troyanskaya et al., 2001] |
| SVDImpute | singular value decomposition | imputation | k = 3 | yes | [Troyanskaya et al., 2001] |
| SVTImpute | singular value thresholding | imputation | $\lambda = 3$ | yes | [Cai et al., 2010] |
| impute.knn | k-nearest neighbor | impute | - | - | [Troyanskaya et al., 2001] |
| cart | classification and regression trees | mice | m = 1 | - | [Doove et al., 2014] |
| mean | arithmetic mean | mice | m = 1 | - | [van Buuren and Groothuis-Oudshoorn, 2011] |
| midastouch | distance aided selection | mice | m = 1 | - | [van Buuren and Groothuis-Oudshoorn, 2011] |
| norm | normal model (linear regression) | mice | m = 1 | - | [van Buuren and Groothuis-Oudshoorn, 2011] |
| pmm | predictive mean matching | mice | m = 1 | - | [van Buuren and Groothuis-Oudshoorn, 2011] |
| rf | random forest | mice | m = 1 | - | [Shah et al., 2014] |
| ri | random indicator | mice | m = 1 | - | [Jolani, 2012] |
| sample | simple random sampling | mice | m = 1 | - | [van Buuren and Groothuis-Oudshoorn, 2011] |
| imputePCA | principal component analysis | missMDA | nboot = 1 | yes | [Josse and Husson, 2012] |
| MIPCA | multiple imputation with pca | missMDA | nboot = 1 | yes | [Audigier et al., 2016] |
| missForest | MV imputation using random forest | missForest | - | yes | [Stekhoven and Buhlmann, 2012] |
| bpca | bayesian pca | pcaMethods | - | yes | [Oba et al., 2003] |
| nipalsPca | nonlinear iterative partial least squares | pcaMethods | - | - | [Wold, 1966] |
| ppca | probabilistic pca | pcaMethods | - | yes | [Tipping and Bishop, 1999] |
| nlpca | nonlinear pca (with neural net) | pcaMethods | - | - | [Scholz et al., 2005] |
| svdImpute | singular value decomposition | pcaMethods | - | - | [Troyanskaya et al., 2001] |
| impSeq | sequential | rrcovNA | - | yes | [Verboven et al., 2007] |
| impSeqRob | robust sequential | rrcovNA | - | - | [Branden and Verboven, 2009] |
| softImpute | soft-thresholded svd | softImpute | - | - | [Mazumder et al., 2010] |
| irmi | iterative robust model-based | VIM | - | yes | [Templ et al., 2011] |

Table 1: Characteristics of the 30 applied imputation algorithms, sorted in alphabetic order by their package name.
